# Exploring Single-Cell Gene Regulatory Dynamics in Rett Syndrome

**DOI:** 10.1101/2025.10.01.679774

**Authors:** Sofia G. Rodriguez, Irene Cartas-Espinel, Camilo Villaman, Mabel Vidal, Eduardo Pérez-Palma, Jesus Espinal Enriquez, Alberto J. Martin, Mauricio A. Saez

## Abstract

Rett syndrome is a monogenic disorder with an incidence of 95% in women, characterized by the complexity of studying the associated phenotype due to the heterogeneity in patient tissues from the stochastic silencing of the affected X chromosome. Furthermore, we are largely unaware of the cascade of alterations that occur in neurons due to transcriptional changes induced by the affected MECP2 gene. To address these challenges, an in-depth network analysis was implemented using organoid single-cell transcriptomic data derived from human patients. We performed a Weighted correlation network analysis and trajectory analysis to understand the differences in the developmental processes between samples, we followed by the generation of gene regulatory networks for each relevant cell developmental pathways to assess the master regulator that are involved in this process, with the differential expressed genes with potential therapeutic implications being identified by integration with SFARI and Genes4Epi. The results were adapted into dynamic Boolean models fitted with the transcriptomic data for validation in which we evaluated the attractor field from each reachable state. These approaches allowed us to explore differences in regulatory behavior in the developmental pathways.

Our study provides an insight that pinpoints the cellular stages on which the regulation and compensatory mechanism activate and regulate Rett syndrome. We identified 19 Master regulators for the Dopaminergic developmental trajectory, as well as 34 Master regulator genes for the Gabaergic developmental trajectory. Dynamic Boolean modeling of these systems showcased a comprehensive understanding of the disrupted developmental pathways of Rett syndrome, highlighting the transitional states of potential within maturation trajectories as the key point of divergence in regulation for Rett syndrome. After complementing with enrichment and clinical relevant variant analysis, we identify the key actors in this system as NR2F1 and TCF4, with TCF4 suggesting a symmetrical compensatory relationship with MeCP2, and NR2F1 as possible link with wider developmental conditions, this concluded with highlighting the possibility of regulation in this condition being affected by the MAPK-ERK pathway of transcriptional regulation, offering a novel angle for targeted research.

## Introduction

Rett Syndrome (RTT) is a rare post-natal neurodevelopmental disorder primarily affecting females[1–3] with 90–95% of diagnosed cases involving the methyl-CpG binding protein 2 (MeCP2), caused by mutations in its corresponding gene (MECP2), located on the Xq28 band of the X chromosome [4,5]. RTT clinical manifestation typically progresses through four stages, initially in early onset RTT patients exhibit a normal development during the first 6-18 months of life, then they experience a rapid deterioration characterized by regression of developmental milestones, motor issues, intellectual disability and autism-like symptoms, this is followed by a plateau stage beginning at approximately 2-10 years old, and maintained during most of the patient life, in which clinical and therapeutic approaches showcase the most success, with the final stage characterized by a progressive deterioration of the patient due to severe motor dysfunction, loss of ambulation, and early mortality due to progressive neuromuscular and autonomic decline [3,6,7].

Despite the following RTT is clinically heterogeneous, with symptom severity and variability of the phenotype being influenced by the location and type of the variant [3,7] as well as by genetic background and cellular environment with truncating variants in MECP2 being associated with an overall higher clinical severity[8]. While some mutations, such as R133C and R306C, affect specific biochemical interactions (e.g., DNA binding or co-repressor recruitment), their clinical consequences can be unexpectedly mild[9–11]. Additionally, the same mutation may yield different phenotypes in different individuals[9,12].

Due to the location of MECP2, the X-chromosome inactivation (XCI) has been proposed to be an important factor in the onset and severity of phenotypic symptoms[2,8,13]. XCI is a stochastic process that takes place in the initial stages of the embryogenesis, causing a mosaic expression of X-linked genes [13–16], in case RTT patients,this results in both MECP2-expressing and MECP2-deficient neurons coexisting[8,13,17]. Despite reported cases of skewed XCI of MECP2 having a protective effect in identical mutations variants over random phenotype[8,10,16],recent studies using allele-specific methylation-based assays have shown that XCI patterns, do not consistently correlate with clinical severity suggesting that multiple regulatory layers beyond XCI contribute to the clinical phenotype [13,17], further illustrating the complexity of the phenotype variations involved in RTT[1,12,18].

MeCP2 is widely recognized as a key modulator of gene expression in neurons, acting through its binding to methylated cytosines in both CpG (mCG) and non-CpG (mCA) contexts[1,4,5,18–22]. It acts primarily as a transcriptional repressor through recruitment of co-repressor complexes such as NCoR/SMRT and histone deacetylase (HDAC) complexes via its transcriptional repression domain (TRD)[5,12], with the most frequent RTT-associated mutations cluster within the segments encoding for the functional domains and disrupting molecular interactions[12,23].

In the methyl-CpG-binding domain (MBD), the T158M mutation reduces DNA binding affinity, while R133C also impairs methylated cytosine recognition[24,25]. The intervening domain (ID) links the MBD and TRD and profoundly influences DNA binding thermodynamics[26], in this region the R168X mutation produces a truncated protein lacking the TRD, abolishing co-repressor recruitment [12,23,25]. Within the TRD, truncating mutations such as R255X and R270X, and the missense mutation R306C disrupt NCoR/SMRT binding, impairing chromatin remodeling and transcriptional repression [12,24,25]. In the C-terminal domain (CTD), L386fs, and E397K, alter histone interactions and activity-dependent phosphorylation events [12,23,24]. Additionally to its canonical function, MECP2 is also implicated in alternative splicing, chromatin compaction, and regulation of non-coding RNAs; with its function extending beyond repression[23] as recent evidence suggest that it can also facilitate transcriptional activation under specific conditions[4,23]. Disruption of these functions results in transcriptional noise, aberrant synaptic maturation, and impaired neural circuit homeostasis, all hallmarks of RTT neuropathology[5,20].

However, despite these insights, the mechanisms that determine whether MeCP2 acts as a repressor or activator remain poorly understood[9,12,23]. In particular, transcriptional regulation alone cannot fully explain the diverse phenotypes observed in Rett syndrome[18,20,23], with recent studies having revealed that MeCP2 occupancy is not limited to methylated DNA[23,25,27]. Chromatin profiling in neurons and mouse brain tissue has identified MeCP2 binding at unmethylated regulatory regions[4,28,29], including promoters[4,29] and enhancers[30], challenging the classical model of MeCP2 as solely a methylation-dependent reader.

These multifaceted and context-dependent interactions point to a broader regulatory landscape than previously appreciated and highlight the current limitations in predicting phenotype from genotype[1,12,18,23] thus understanding how MECP2 operates across diverse cellular and molecular contexts will be essential for linking its molecular interactions to the complex phenotypes of RTT[5,12,18].

Given the molecular and cellular heterogeneity in RTT, bulk transcriptomic approaches are insufficient to resolve the full scope of transcriptional dysregulation [23,31–33]. These methods average gene expression across mixed populations of cells, thereby masking the diverse effects of MECP2 mutations[31,34]. To overcome this limitation, single-cell RNA sequencing (scRNA-seq) allows the resolution of gene expression at the level of individual cells, uncovering cell-type-specific transcriptional signatures and subtle differences that would otherwise be obscured[35,36].

Given the cell-type-specific heterogeneity captured by scRNA-seq, and the intrinsic complexity to of MeCP2 we require a framework capable of encompassing and embracing the complexity of RTT phenotype[36–40], for this we decided that biological network analysis can allow us to model, explore and understand the cascade of MECP2[39,41,42], as the analysis of condition specific networks can both preserve the complexity within the system and reduce to a way that allows us to understand key mechanisms within the condition[38–41].

We employ Weighted Gene Correlation Network Analysis (WGCNA) to construct We employ Weighted Gene Correlation Network Analysis (WGCNA) to construct correlation networks by clustering genes that exhibit similar expression patterns across individual cells, allowing the detection of coordinated transcriptional programs associated with specific cell types or states, obtaining group genes with coordinated expression patterns into biologically meaningful modules[43,44]. These modules are clusters of highly interconnected genes, corresponding to clusters of genes with high absolute correlations and summarized by highly connected genes that become representatives of the gene expression profiles of the module(eigengenes)[44,45].Complimenting these eigengenes by utilizing functional enrichment, offering insight into the biological roles of distinct gene sets revealing interactions between cellular processes or transitions between cell states, which could allow us to identify transcriptional signatures associated with MECP2 dysfunction that may not be directly inferred through regulatory logic but reflect shared cellular phenotypes or stress responses within the patient[44,46,47].

However this method alone allows us to evaluate possible correlations, it still doesn’t capture the regulatory landscape in RTT. To explore these processes we apply gene regulatory network (GRN) inference, which captures directed relationships between transcription factor (TF) and their target genes[42,48]. This systems-level approach coils the identification of regulatory hubs and key nodes that control disease-relevant expression patterns[49,50]. In the context of RTT, where MECP2 acts as a global transcriptional modulator, GRNs offer a critical insight into both direct and downstream effects of MECP2 dysfunction showcasing possible compensatory mechanisms[51] and cell-type-specific regulatory dynamics[52] that may not be apparent in linear gene expression analyses.

To capture the temporal behavior of these regulatory circuits, we extend our analysis by integrating GRNs with dynamic Boolean modeling. Boolean models represent gene activity in binary states (ON/OFF) and simulate how regulatory interactions evolve over time, based on logical rules derived from network topology and expression data[53–56]. This allows us to explore how MECP2-related perturbations propagate through the network, revealing dynamic shifts in regulatory state[56], trajectory-specific vulnerabilities[54], and potential windows for therapeutic intervention. Unlike static network analyses, Boolean modeling incorporates the dynamic nature of gene regulation, offering predictive insights into disease progression and enabling in silico testing of regulatory hypotheses

By combining scRNA-seq, WGCNA, GRN inference, and dynamic modeling, our approach provides a comprehensive framework to dissect the regulatory consequences of MECP2 dysfunction in RTT. This integrative strategy allows for the identification of robust and biologically relevant targets by evaluating their regulatory centrality, dynamic behavior, and cell-type specificity—ultimately supporting the development of more effective, mechanistically informed therapeutic intervention

## Methods

### Data

The dataset GSE165577, generated by Samarasinghe et. al[57]. was obtained from the raw BCL file obtained from the Gene Expression Omnibus (GEO)[58].The dataset contained scRNA-seq data from cerebral organoids generated from isogenic control (WT) or MECP2 mutant (705delG/ E235fs; Mut) induced pluripotent stem cells were derived from a Rett syndrome patient, and collected at 3 time points: 56 days (8w), 70 days (10w), and 100 days (10w). Day 56 pools each contain 3 unfused Cortex and 3 unfused Ganglionic eminence organoids. Day 70 and 100 pools contain 3 Cortex+Ganglionic eminence fusion organoids control (WT) condition and the affected (MECP2-) condition, with an experimental window of incubation of the samples of 50 days (8w), 76 days (10w) and 100 days (14w) with 2 biological reproduction per harvest. The test platform used was a GPL11154 Illumina HiSeq 2000 (Homo sapiens).

### ScRNA-Seq Data Processing

Then the data was aligned using GRCh38.p14 as the reference genome with 10x Genomics Cell Ranger (v 8.0.0)[59] in its default settings, for all samples and reproductions. The resulting count matrix files were exported and analyzed into Seurat (v5)[60] creating 3 main analysis seurat objects, the WT seurat object composed of all the control samples in all ages, the MECP2-seurat object composed of all the affected samples, and Combined seurat object composed of all samples, to facilitate some of the further analysis; with Age being kept in the metadata ob each individual object.

Quality control measures included filtering out cells expressing fewer than 200 genes and genes detected in fewer than 3 cells. Cells exhibiting over 5% mitochondrial gene expression were excluded to remove potential apoptotic cells. Before preprocessing, the dataset contained null cells and null genes. After filtering, null cells and null genes remained, ensuring a high-quality dataset for analysis.Data normalization was performed using the LogNormalize method, scaling gene expression to 10,000 transcripts per cell. Highly variable genes were identified using the seurat method, selecting the top 2000 variable genes for downstream analysis.

Principal Component Analysis (PCA) was conducted on the scaled data, retaining the top 15 principal components. A nearest-neighbor graph was constructed using the 15 nearest neighbors. Clustering was performed using the Leiden algorithm with a resolution parameter of 1. Dimensionality reduction for visualization was achieved using Uniform Manifold Approximation and Projection (UMAP).

Differential expression analysis was conducted to identify marker genes for each cluster. The Wilcoxon test was employed to determine statistical significance, with a threshold of adjusted p-value < 0.05. The top 3 differentially expressed genes per cluster are available in Annex1.

### Cellular Annotation

Automated annotation of cell types was performed using the ScType framework. We applied ScType v.3.0 [61] to the RNA assay of the integrated Seurat object, using tissue-specific marker sets from the Brain ScTypeDB. ScType assigns cell types based on the expression of positive and negative gene signatures, computing a score for each cell type within each cluster. For each cluster, the top-scoring cell type was selected, and low-confidence assignments—defined as scores lower than one-quarter of the number of cells in the cluster—were labeled as “Unknown”.

Manual validation was performed by reviewing the expression of established marker genes to confirm consistency between assigned cell types and expected expression profiles. The resulting annotations were added to the Seurat object metadata and visualized using UMAP, grouped by inferred cell type.

During the Trajectory Analysis some cells belonging to the Unknown group were re-classified due to the reduction process, these cells were retroactively flagged for population distribution evaluation of cell types in the samples.

### Weighted Correlation Network Analysis

WGCNA was performed on the processed Seurat objects generated as described in the *ScRNA-Seq Data Processing* section, from these the Radial Glial cells and Gabaergic neurons were the subsets chosen for analysis. We utilized the hdWGCNA (v 0.4.06)[62] in its R protocol analysis, genes expressed in at least 5% of cells were selected. Metacells were generated using *Age* and *Condition* as grouping variables for the cell type subset, with a k-nearest neighbor approach (*k* = 25). These metacells were normalized and transformed into the required WGCNA input format.

The soft-thresholding power was determined, and the resulting table was exported. The weighted gene co-expression network was constructed using a soft power of 9. Module eigengenes were calculated for each group defined by *orig.ident*, and both harmonized and non-harmonized module eigengenes were obtained. Module connectivity was assessed to identify hub genes, and module expression scores were calculated using the top 25 genes per module according to the Seurat method, resulting in further outputs available in Annex 2.

### Gene Set Enrichment Analysis

Gene set enrichment analysis was performed using the R package enrichR[63] with the Enrichr online database. The connection to the Enrichr server was established, and three databases were selected for analysis: GO_Molecular_Function_2021, GO_Biological_Process_2021, and KEGG_2021_Human. A list of genes obtained from Cytoscape[64] was imported and converted into the character vector format required by enrichR. The enrichment query was run against the selected databases, generating separate result tables for each.

For downstream analysis, results from each database were filtered to retain only terms with *p*-value ≤ 0.5. The filtered GO Molecular Function and GO Biological Process results were combined into a single table. Visualization was performed using `plotEnrich`, generating bar plots of the top 5 enriched terms (truncated to 40 characters) for GO and KEGG pathways, ordered by *p*-value. The final filtered GO and KEGG results were exported as CSV files for further inspection.

To complement the enrichment results, disease association analysis was performed using the Orphanet disease–gene database. The Orphanet dataset was imported as a CSV file, and the gene list for each master regulator was loaded from a text file containing one symbol per line. Each disease entry in the Orphanet table was parsed to retain only genes present in the input list, and rows with at least one matching gene were selected. The *Genes* column was updated to display only the overlapping symbols, and the resulting table of matched diseases and their associated genes was exported as a CSV file Annex. 3.

### Pseudotime Trajectories

Pseudotime trajectory inference was performed using Monocle 3[65], following the workflow outlined in the *Calculating Trajectories with Monocle 3 and Seurat vignette*. Low-quality cells were excluded based on mitochondrial content (<10%) and RNA feature thresholds (nFeature_RNA > 500 and below mean + 3×SD). The filtered object was split by sample (`orig.ident`), normalized, and processed to identify variable features. After selection of integration features, each subset was scaled and subjected to PCA. Integration was performed using reciprocal PCA (RPCA) via `FindIntegrationAnchors`, followed by `IntegrateData`.

The resulting integrated object was scaled, reduced with PCA and UMAP, and clustered. It was then converted into a Monocle 3 `cell_data_set` using `as.cell_data_set`. Cells were clustered with `cluster_cells`, and the main partition (partition 1) was selected to ensure a coherent trajectory structure. A new CDS object was created from this subset, and trajectory learning was performed using `learn_graph`. The resulting graph was saved and used for downstream pseudotime analysis and visualization.

### Inference of gene regulatory networks

To prepare expression matrices for gene regulatory network inference, we first excluded non-neuronal cell types, such as cancer and immune system cells, from the full dataset. To generate dopaminergic and GABAergic neuron-specific subsets, we filtered out clusters and cell types corresponding to each other’s lineage. The resulting subsets were visualized using UMAP by both cell type and Monocle3 cluster identity. To ensure the full gene landscape of the cells within the monocle3 objects, we matched cell barcodes with the full dataset and updated metadata accordingly. Both Dopa and Gaba objects were then merged using JoinLayers to unify the RNA counts and normalized with the LogNormalize method. Finally, we generated count matrices for each Monocle3 cluster and cell type, stratified by experimental condition. Each matrix was transposed and exported as a comma-separated file for downstream network inference

We inferred a gene regulatory network using Arboreto’s implementation of GRNBoost2[66]. To this end, we loaded the expression matrix (GABAergic neurons_MECP2-Gaba.txt) as a pandas DataFrame, with gene names as column headers. Transcription factor (TF) names were imported using Arboreto’s load_tf_names utility function. We then ran the GRNBoost2 algorithm (arboreto.algo.grnboost2) with the expression matrix and the TF list to infer TF-target gene interactions. The resulting network was saved as a tab-separated file without headers or row indices.

Subsequently, we filtered the inferred network to retain the top 10% of target gene connections for each TF. To do this, we first extracted all unique TFs (source genes) from the network file. For each TF, we identified all corresponding TF-target gene pairs, sorted them by their importance score (“value”), and selected the top decile of interactions. These filtered interactions were then printed in a tab-separated format for downstream analysis Annex. 4.

### Inference of master regulators

Gene regulatory networks were visualized using Cytoscape (v3.10.2) [64]. To evaluate the biological relevance of the predicted interactions, we compared them to high-confidence regulatory interactions from TF-link, a human-specific resource that integrates curated and experimentally supported TF-target relationships from multiple databases.

We began by intersecting a predefined list of transcription factors with the gene expression matrix to retain only TFs present in the dataset. In order to focus on transcription factors that are both expressed and condition-relevant when comparing our inferred networks to the TF-link reference.

We identified candidate master regulators by selecting the first and second upstream neighbors of MECP2 and ranking transcription factors by their out-degree within the filtered network, following the topological inference of master regulators (MRs) described by Davis and Rebay [67]. We iteratively removed nodes with the lowest out-degree until further removal disconnected the network. All curated GRNs and identified master regulators were visualized and analyzed in Cytoscape, and the corresponding session files are available for further inspection Annex. 5.

### Differential gene expression analysis with integration with clinically relevant databases

For the genetic burden analyses, we generated gene lists for each of the cell types and the Monocle3 inferred pseudotimes, to increase the possible pool of solutions these were obtained from the total matrix objects including all identified cell types and all monocle3 groups. These lists were tested for enrichment against established phenotype-related gene sets, including epilepsy and autism. Specifically, we used the Genes4Epilepsy [70] database for epilepsy and the SFARI database [71] for autism.

To test whether the overlap between the generated lists and the phenotype-related databases exceeded random expectation, we applied a hypergeometric test. This test is appropriate in this context, as it models sampling without replacement from a finite population (the genome). For each comparison, we calculated the probability of observing at least K overlapping genes under the null hypothesis and reported the P-value, odds ratio, and number of overlapping genes.

To establish an empirical baseline, we repeated the analysis 1000 times using randomly generated gene lists of equal size to the original sets. The distribution of P-values from these random repeats was compared to the observed P-value to obtain an adjusted enrichment measure. In total, 1052 genes from the Genes4Epilepsy database and 1238 genes from the SFARI database were included in the comparisons.

Subsequently, both databases were filtered for clinically relevant genes using the ClinGen Gene-Disease Validity resource [72],retaining only genes with a supported gene–disease validity link. This filtering resulted in 671 genes from the Genes4Epilepsy database and 480 genes from the SFARI database.

### Dynamic Boolean Modeling

We applied a combination of Boolnets[68] and BoNesis[69] to infer Boolean networks that are consistent with both prior regulatory knowledge and observed gene expression dynamics. The influence graph topology was constructed using the previously inferred gene regulatory network.

For the observational component, we determined the nature of each regulatory relationship (+/–) based on the correlation between regulator–target pairs, focused on the WT and MECP2-conditions in separated evaluation of the systems. For the development lineage Cells were subset from each respective Seurat object by condition, and the lineage-MRs were loaded as an external gene list. All pairwise combinations of MRs were assessed using Pearson correlation based on the expression levels across cells. These were calculated and visualized using `FeatureScatter` plots in seurat, grouped by cell type to assess the co-expression relationships among regulators and support the relationship in the GNR.

To convert the expression data into a format suitable for Boolean modeling, we performed cluster-level binarization using the BoolNets package in R. For each condition, cells were grouped by pseudotemporal clusters, and average expression profiles were computed with Seurat’s `AverageExpression`. The resulting matrices were extracted with `GetAssayData` and binarized using the `binarizeTimeSeries` function with the `edgeDetector` method. The final binarized matrices were transposed and exported.

The Boolean modeling was conducted using BoNesis in python. This information served as the basis for the synthesis of Boolean networks whose structure is consistent with the prior influence graph and whose dynamics reproduce the transitions observed across cell states in the binarized matrix. The dynamical constraints were defined by default of the library, encoding each cell type as a steady state and formulating reachability constraints between them under the Most Permissive Semantics.

BoNesis was used to enumerate up to 5,000 Boolean networks that satisfied these structural and dynamical constraints. Attractors were computed as minimal trap spaces of the Boolean network and represented as hypercubes mapping every network component to 0, 1, or *. Static attractors corresponded to fixed points with stable gene activation patterns, whereas cyclic attractors represented recurrent regulatory sequences. For each analysis, we used the transcriptional state of a given cell type as the `reachable_from` condition, thereby restricting the attractor search to trajectories accessible from that state.

To analyze the dynamical behavior of the models, we computed the attractors landscape reachable from each defined starting state corresponding to major cell populations from the developmental line. For each initial state, we collected attractors across all models and determined the frequency and basin size for each attractor configurations. This was done by aligning the attractors across models, identifying common patterns, and extracting consensus profiles. The resulting attractor sets represent stable or recurrent transcriptional configurations associated with distinct stages of differentiation. We compared attractor landscapes between wild-type (WT) and MECP2– models constructing the combined universe of attractors across both conditions. It then evaluates which attractors are preserved, lost, or gained in MECP2– compared to WT, while also quantifying their similarity using Hamming distances, estimating attractor frequencies and basin sizes to assess their relative stability and prevalence.

## Results

### Results 1: Analysis of single cell samples reveal differences in population distribution between Rett syndrome and control samples

In order to evaluate the distribution of cell type population between control samples (WT), and RTT affected samples (MECP2-), we performed a SC transcriptomic data analysis, utilizing 10x Chromium Cellranger for quality control, alignment, and then proceeding further analysis with Seurat (v5) to explore the cells properties and distribution. PCA, Jackstraw, and UMAP were used for clustering and dimensionality reduction to identify relevant cell populations. The Cell-type annotation was initially performed automatically using the Sc-Type software, and cells were subsequently verified through manual review by examining representative cellular markers. This analysis was performed for WT (Fig1 .A) and MECP2-(Fig.1. B) global conditions at 8w, 10w, and 14w.

**Figure 1:**
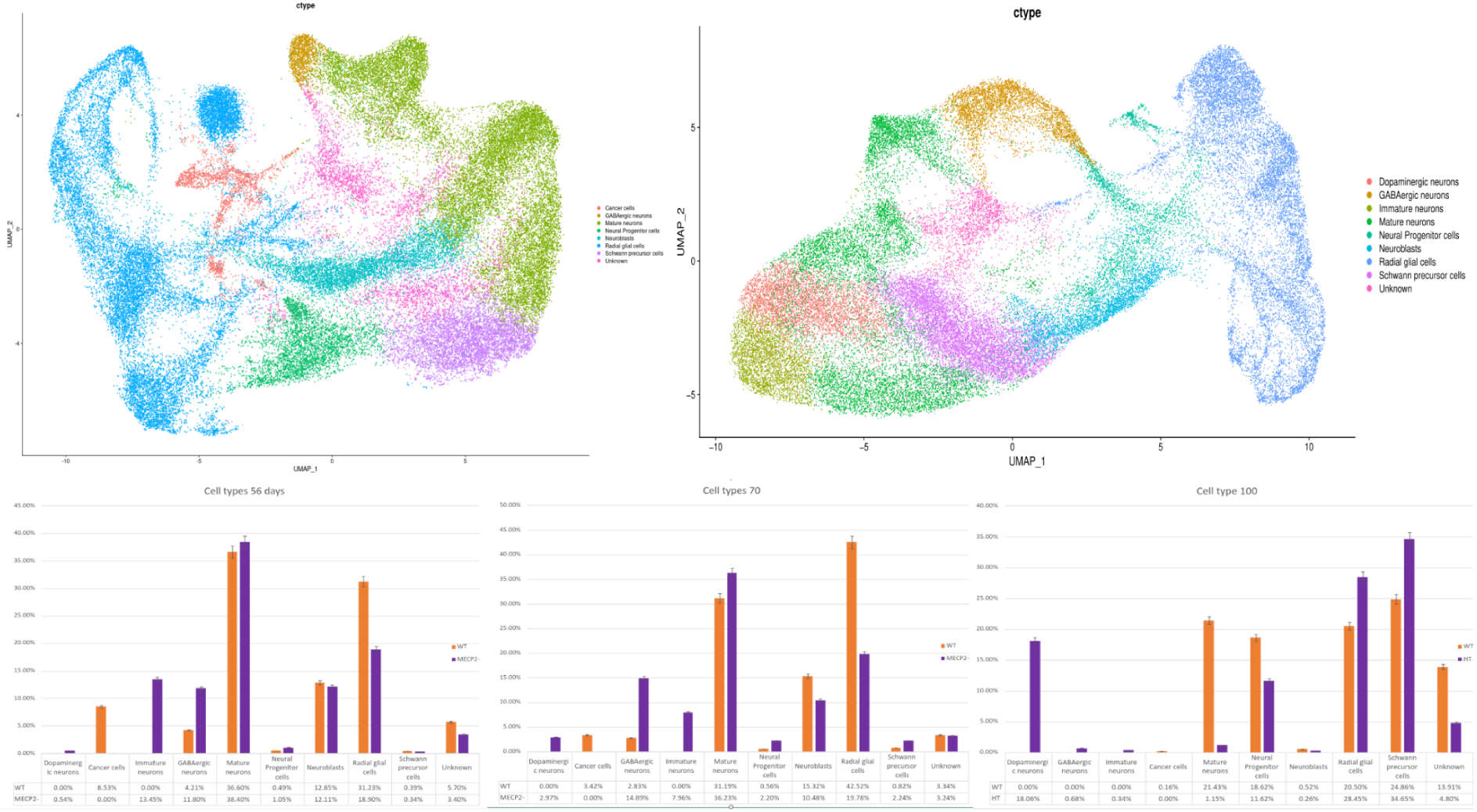
A) 1) UMAP showcasing the different cell types identified within the WT condition B.) UMAP showcasing the different cell types identified within the MECP2-condition. C) 1) bar graph showcasing the distribution of identified populations between WT (orange) and MECP2-(purple) at 8 weeks alongside the population distribution table per condition. 2) bar graph showcasing the distribution of identified populations between WT (orange) and MECP2-condition(purple) at 10 weeks. 3) bar graph showcasing the distribution of identified populations between WT (orange) and MECP2-condition(purple) at 14 weeks.

Sc-Type classified cells into ten populations, corresponding to the following distribution in the WT condition: GABAergic neurons (2.09%), Mature neurons (28.84%), Immature neurons(3.57%), Neural progenitors (7.74%), Neuroblasts (8.72%), Radial glial cells (30.53%), Schwann cell progenitors (10.28%), and others cell types (8.22%), which constitute a group of minority cell populations corresponding to less than 0.5% of the cell populations and non-neuronal cell types (see Table1.). In the MECP2-condition, the cell populations consisted of Dopaminergic neurons (9.33%), GABAergic neurons (7.25%), Mature neurons (20,39%), Immature Neurons (6,06%), Neuronal progenitors (6,28%), Schwann cell progenitors (16,91%), Neuroblasts (6,17%), Radial glial cells (23,59%), and other cell types (4.09%) (see Table1.).

From this analysis the most significant insight is the distinct variation of Dopaminergic cells between WT and MECP2-, where less than 0.5% of the total cell population in WT and MECP2-corresponds to 9.33% of the total population, respectively. Similarly it is possible to observe changes in the distribution between WT Radial glial cells total corresponding to 30.53% of the total cell population and MECP2-corresponding to 23.59% of the total population.

**Table 1:** Table describing the showcasing the population percentage per condition.

| Cell types | WT | MECP2- |
| --- | --- | --- |
| GABAergic neurons | 2.09% | 7.25% |
| Dopaminergic neurons | < 0.5% | 9.33% |
| Mature neurons | 28.84% | 20.39% |
| Immature neurons | 3.57% | 6.06% |
| Neural progenitors | 7.74% | 6.28% |
| Neuroblasts | 8.72% | 6.17% |
| Radial Glial Cells | 30.53% | 23.59% |
| Schwann cell progenitors | 10.28% | 16.91% |
| Other cell types | 8.22% | 4.09% |

In a deeper insight we evaluated the differences across the population at the different time points with the population of Dopaminergic neurons at 8w representing 0.54% of the total population(Fig 1. C-1), at 10w representing 2.97% of the total cells(Fig 1. C-2) and at 14w representing 18.06% of the total cells (Fig 1. C-3).

This is accompanied by a difference in the distribution of neural precursor cells and undifferentiated neurons in the samples, with a reduction rate over the time consistent between both conditions. Neural precursor Cells having a 0.1% - 0.5 % increase during 8w and 10w(Fig 1. C-1-2), having a 32x increase by 14w in the WT population (Fig 1. C-3), similarly MECP2-condition had a 0.5 - 2% increase in population in 8w and 10w (Fig 1. C-1-2), and 4x increase in 14w (Fig 1. C-3), with both presenting a low increase up to week 14 with an explosive increase of higher magnitude in the control condition. Similarly in Schwann Precursor Cells, with a 1.11% increase during 8w and 10w (Fig 1. C-1-2), having a 29x increase by 14w (Fig 1. C-3) in the WT population, and, the MECP2-condition presenting a 5% increase in population in 8w and 10w(Fig 1. C-1-2), and 14x increase in 14w(Fig 1. C-3).

However in the case of Radial Glial cells, we have a significant difference in behavior overtime between condition, for the Wt condition we have a slight 0.36% growth between 8w and 10w, by 14w there is decrease of a 96% of the population; however for MECP2 condition between 8-10w there is a non relevant growth (<0.5%), but by 14w we have a 43.8% increase in population.

Finally in the reduced population, cancer cells were identified as germinal cells for neurons, as their changes in this population complement the influx of precursors seen at 14w (Fig 1. C-3). The population at 8w with a difference of 12.32% between conditions (Fig 1. C-1), at 10w representing 22.74% of the total (Fig 1. C-2) and at 14w representing 7.95% between the conditions (Fig 1. C-3).

This allowed us to appreciate key differences between the distribution and changes of the population over time, revealing distinct differences in cell type distribution between WT and MECP2-conditions. Dopaminergic neurons, which constitute less than 0.5% of the WT population, are expanded to 9.33% in MECP2-samples, with a progressive increase over time. Neural precursor and Schwann cell progenitor populations show a greater increase at 14 weeks in WT compared to MECP2-. In contrast, radial glial cells exhibit opposite trends, with a sharp decline in WT at 14 weeks, whereas in MECP2-, the population increases. These findings indicate significant alterations in cell population dynamics between the two conditions over developmental time points.

### Results2: WGCNA analysis reveals differences in maturation between WT and MECP2-cells, in key cellular subtypes

WGCNA was performed to find possible gene modules that could provide clinical interest. For this, we combined WT and MECP2-data, and performed the analysis based on cell types that could provide possible insights for therapeutic targets, while preserving the labels obtained in the previous SC analysis. We aimed to assess whether the differences between WT and MECP2-were significant enough to distinguish them in an unsupervised manner. We utilized Weighted Correlation Networks from the hdWGCNA library in R, which is optimized for high density data, and is able to directly use Seurat objects with expression information (see. Fig 2). Complementary, the criteria of “cell type of interest” was defined as the cells that could provide insights in application in diagnosed patients over cell types that could provide developmental insights. Finally, GABAergic neurons and Radial glial cells were analyzed.

**Figure 2:**
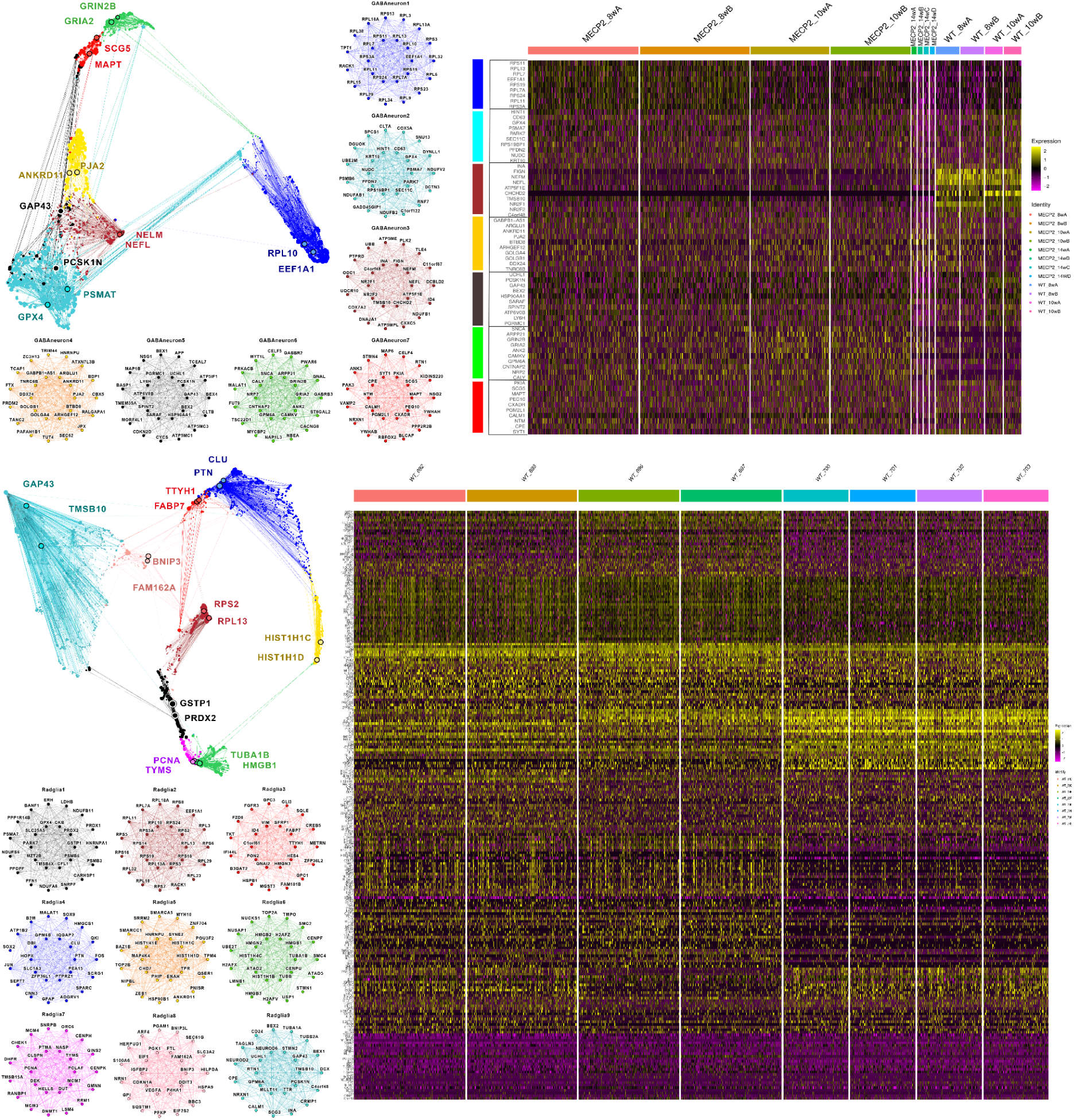
A) UMAP of modules and population of cells per module. B to H) Gene correlation network of the identifying centroids for the corresponding population, indicating in the central circle the 10 most significant genes for the population, surrounded by the 15 secondary ones. I) Heatmap indicating the expression of the modules within the samples ages of the cell type. J) UMAP of modules and population of cells per module.K to S) Gene correlation network of the identifying centroids for the corresponding population, indicating in the central circle the 10 most significant genes for the population, surrounded by the 15 secondary ones. T) Heatmap indicating the expression of the modules within the sample ages of the cell type.

Based on these analyses, it was possible to obtain groups of genes co-expressed (Module) by defining a specific behavior in a population of cells within the cell type. The 25 most representative co-expressed genes per group were designated as centroids, which are the genes that best characterize the behavior and properties of each group.

The modules were dimensionalized in a UMAP containing all the identified modules per cell type (fig with WGCNA Umaps), with the centroids for each found module showcased as a circular network with the inner circle showcasing the top 10 most defining genes per module, this was also complemented with the functional enrichment of the centroids using EnrichR in order to understand their current function and the behavior in the population of cells. Finally, The top 10 genes per module in each cell type over the time points of interest were selected to understand the behaviour during development and their possible contribution into the phenotype.

We began with the Gabaergic neurons. It was possible to observe that the major cellular contribution came from the samples of 8w and 10w. For this cell type it was possible to identify 7 modules with their respective centroids and characteristic expression profiles (see fig. WCGab. A.). The genes associated with module 1 showed higher expression at 8w in MECP2-than WT samples. However, for the 10w period, the expression was similar in both conditions. In module 1 the proteins are particularly associated with protein synthesis processes in cytoplasm (see WCGab B).

Genes associated with modules 4 and 5 are mainly present in the WT samples, with a reduced expression at the 8 weeks MECP2-samples, but was restored at 10 weeks at similar expression levels to the 8 weeks WT. The module 4 centroids are particularly associated with RNA binding and regulation of the processing of metabolic macromolecules (see Fig. WCGab. E); Module 5 centroids are particularly associated with the assembly of cellular protein complexes (see Fig. WCGab. F).

Module 6 is mainly expressed in the MECP2-samples, particularly at 10 weeks of incubation. However, it had the same expression pattern in the WT condition at 8 weeks and 10 weeks. Genes associated with module 7 are consistently expressed in the WT condition, however, its reduced expression in the 8 weeks MECP2-condition and absence in further samples; their centroids are particularly associated with trans-synaptic anterograde signaling (see Fig. WCGab. G) and centroids for module 7 are particularly associated with synaptic processes (see WCGab. H). Extensive list of genes by modules and their score is disponible in supplementary table Anx.5.

In order to evaluate how the identified modules are expressed over time in the sample, we visualized the 10 central genes of each centroid in a heatmap with the Y axis representing the genes and modules, and the X axis showcasing the age of the samples within each cell type. From this analysis we were able to evaluate WT and MECP2-conditions, revealing distinct temporal expression patterns across the cell types of interest some gene modules showed reduced expression in MECP2-at early time points but normalized by 10 weeks, while others were consistently reduced or absent in MECP2-. In radial glial cells, gene modules associated with transcriptional regulation, cell proliferation, and neuronal function exhibited different expression trajectories between WT and MECP2-, with some modules showing delayed or absent expression in MECP2-.

In Radial Glial Cells, the highest population density was observed in the WT samples at 8 and 10 weeks, with a lower presence at 14 weeks, which accounts for the maturation processes in the organoid, and also a lower presence at 14 weeks in all MECP2-samples. For this cell type, nine modules with their respective centroids and characteristic expression profiles were identified (see Fig.2 J).

The genes associated with module 1 were mainly expressed in the WT condition at 10w. TTR showed sporadic expression at 8w, a significant increase at 10w, and a decrease at 14w in WT. In contrast, in MECP2-, it was sporadically expressed at 8w and 14w but downregulated at 10w. The centroids of module 1 are particularly associated with the regulation of planar polarity (see Fig.2 K).

The genes associated with module 2 were predominantly present in the 8w and 14w WT conditions, while in MECP2-, they were sporadically and negatively expressed. The centroids of module 2 are particularly associated with the catabolic process of nuclear transcription of mRNA (see Fig.2 L).

The genes associated with module 3 were expressed at low levels in 8w WT, sporadically in 10w WT, and were downregulated at 14w WT. However, in MECP2-, the 8w and 10w samples showed expression patterns similar to 8w WT, whereas the 14w samples resembled 10w WT. The centroids of module 3 are particularly associated with the negative regulation of transcription by RNA polymerase II and the regulation of cell proliferation (see Fig.2 M).

The genes associated with module 4 were predominantly expressed in 14w WT, while in MECP2-, their expression was downregulated at the same stage. In both conditions, they showed sporadic expression in the 8w and 10w groups. The centroids of module 4 are particularly associated with cellular response to cytokines and negative regulation of cell proliferation (see Fig.2 N).

The genes associated with module 5 were mostly expressed in the 10w MECP2-condition. However, in the WT condition, their expression was higher in the 8w samples and gradually decreased in the 10w and 14w samples. The centroids of module 5 are particularly associated with transcriptional regulation by RNA polymerase II (see Fig.2 O).

The genes associated with module 6 were particularly expressed in both WT and MECP2-conditions at 14w, with sporadic expression in the other time groups. Notably, HCPX and HST1H1D were expressed in opposite patterns between WT and MECP2-. The centroids of module 6 are particularly associated with RNA binding and interaction with DNA (see Fig.2 P).

The genes associated with module 7 were particularly involved in DNA metabolic processes (see Fig.2 Q).

The genes associated with module 8 were mainly expressed in 8w samples, with their expression significantly decreasing in 10w WT and 14w MECP2-groups, maintaining similar expression patterns between conditions. The centroids of module 8 are particularly associated with apoptosis and hypoxia processes (see Fig.2 R).

Finally, the genes associated with module 9 were consistently expressed in 8w WT and 8w-10w MECP2-, but their expression significantly decreased by 14w in MECP2-. The centroids of module 9 are particularly associated with nervous system development and neuron generation (see Fig.2 S).

In order to evaluate how the identified modules are expressed over time in the sample, we visualized the 10 central genes of each centroid in a heatmap with the Y axis representing the genes and modules, and the X axis showcasing the age of the samples within each cell type. From this analysis we were able to evaluate WT and MECP2-conditions, revealing distinct temporal expression patterns across the cell types of interest some gene modules showed reduced expression in MECP2-at early time points but normalized overtime, while others were consistently reduced or absent in MECP2-. In radial glial cells, gene modules associated with transcriptional regulation, cell proliferation, and neuronal function exhibited different expression trajectories between WT and MECP2-, with some modules showing delayed or absent expression in MECP2-however overall behavior seems related to development rather than distinct MECP2-effects.

These results indicate that gene co-expression patterns in these cell types follow different developmental trajectories in MECP2-compared to WT, suggesting changes in the coordination of the processes over time.

### Results 3: Analysis of GRN reveals a core system of MECP2 master regulators shaping neuronal developmental trajectories and phenotype control

In order to find possible new therapeutic targets for RS, a GRN analysis was performed. We performed a cell-type annotation using ScType and manual validation with corresponding cell markers to the monocle3 generated cell types developmental lines. For these processes we focus on the search of master regulators within the developmental pseudotime trajectories, to explore key regulatory components between the networks, for this we generated GRN per cell type and condition for the Gabaergic development and the Dopaminergic Development; obtaining MECP2 main neighbors (2nd Neighbors) and uniting them to obtain the Master regulators in the developmental process, we contrasted the found relationships with reported data from the TF-Link database as reference to solidify the finding in this section.

From the dopaminergic neuron development trajectory we identified 1358 transcription factors in the samples of all cell types. Among these, 10 were classified as MECP2 MR in the WT condition and 9 in the MECP2-condition, with MEIS2 being shared between both conditions (see Fig 3. A).

**Figure 3:**
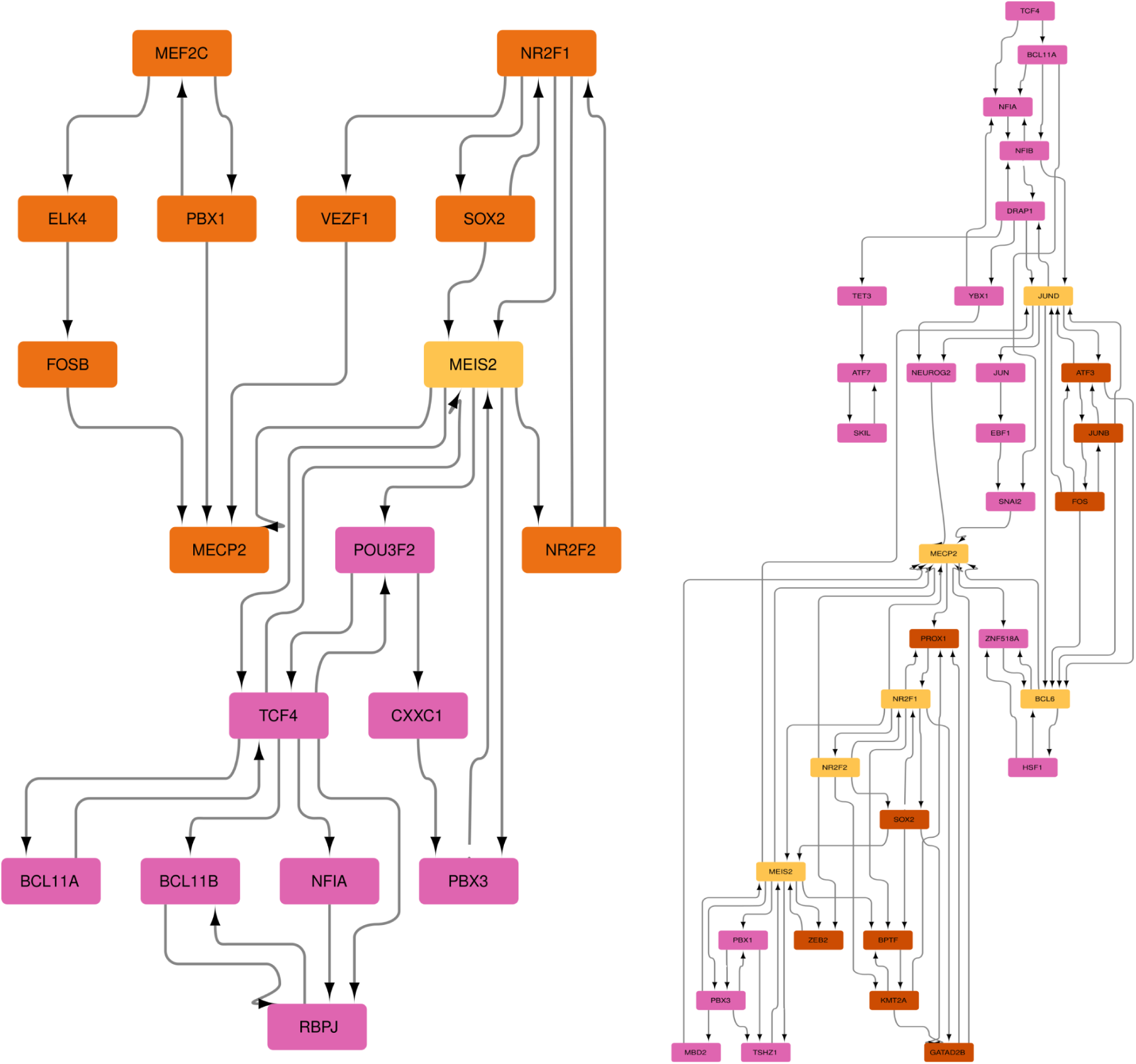
A) Dopaminergic Trajectory Master Regulator Network, with WT condition MR in color orange, common Mr in yellow and MECP2-condition Mr in purple B) Gabaergic Trajectory Master Regulator Network, with WT condition MR in color orange, common MR in yellow and MECP2-condition MR in purple.

In the WT condition, we identified 9 MRs: FOSB, NR2F2, ELK4, MECP2, PBX1, NR2F1, MEIS2, MEF2C, SOX2, and VEZF1. Functional enrichment indicated the Biological Process to be “Regulation of DNA templated Transcription” (GO:0006355), alongside its molecular function being defined as “RNA Polymerase II transcription regulatory region sequence specific DNA binding” (GO:000977), a more detailed enrichment utilizing GeneCards, corroborated this information and finding of particular interest ELK4, as a transcriptional repressors with established roles in chromatin remodeling and early differentiation,and NR2F2 and PBX1 role as developmental regulators with broad functions in organogenesis. Additional enrichment with the Orphanet Database linked their dysfunction with other developmental disorders (see Anex.GRN1), with the most relevant matches indicating that dysfunctions in these systems had a recurrent impact in cardiac development, and intellectual disability.

In the MECP2-condition, we identified 8 unique MRs: TCF4, PBX3, CXXC1, POU3F2, BCL11B, BCL11A, RBPJ, NFIA. Functional enrichment indicated the Biological Process to be “Regulation of DNA templated Transcription” (GO:0006355), alongside its molecular function being defined as “RNA Polymerase II transcription regulatory region sequence specific DNA binding” (GO:000977), a more detailed enrichment utilizing GeneCards, corroborates this highlighting particularly CXXC1 a is involved in epigenetic regulation and nucleosome remodeling and BCL11A as a non-coding regulator implicated in chromatin architecture and long-range gene silencing. Additional enrichment with the Orphanet Database linked their dysfunction with other developmental disorders (see Anex.GRN1), with the most relevant matches being TCF4 associated with Pitt-Hopkins syndrome and YY1 association with Gabriele-de Vries syndrome. BCL11A and ZNF263 are non-coding regulators implicated in chromatin architecture and long-range gene silencing.

From the GABAergic neurons development trajectory we identified 1366 transcription factors. Among these, 15 were classified as master regulators (MRs) in the WT condition and 25 in the MECP2-condition with 6 MRs were shared between both conditions: NR2F1, MEIS2, NR2F2, JUND, BCL6 and MECP2 (see Fig 3. B). Functional enrichment indicated the Biological Process to be “Regulation of DNA templated Transcription” (GO:0006355), alongside its molecular function being defined as “RNA Polymerase II transcription regulatory region sequence specific DNA binding” (GO:000977), a more detailed enrichment utilizing GeneCards and orphanet corroborates this highlighting MEIS2 and NR2F1 as association with intellectual disability.

In the WT condition, we identified 9 unique MRs: JUNB, KMT2A, FOS, ZEB2, ATF3, SOX2, GATAD2B, BPTF and PROX1. Functional enrichment indicated the Biological Process to be “Regulation of DNA templated Transcription” (GO:0006355), alongside its molecular function being defined as “RNA Polymerase II transcription regulatory region sequence specific DNA binding” (GO:000977), a more detailed enrichment utilizing GeneCards and Orphanet corroborates this highlighting KMT2A as linked to Wiedemann-Steiner syndrome and chromatin remodeling during neurodevelopment, with ZEB2 being associated with Mowat-Wilson syndrome, with FOS and JUNB being immediate early-response genes involved in activity-dependent transcription and synaptic plasticity.

In the MECP2-condition, 19 unique MRs were identified: DRAP1, JUN, NEUROG2, TSHZ1, NFIB, YBX1, NFIA, HSF1, TET3, PBX1, ZNF518A, PBX3, EBF1, TCF4, SKIL, SNAI2, ATF7, BCL11A and MBD2. MFunctional enrichment indicated the Biological Process to be “Regulation of DNA templated Transcription” (GO:0006355), alongside its molecular function being defined as “RNA Polymerase II transcription regulatory region sequence specific DNA binding” (GO:000977), a more detailed enrichment utilizing GeneCards and Orphanet corroborates this highlighting MBD2 and TET3 involvement in chromatin accessibility, DNA demethylation, and transcriptional repression with TET3, in particular, being linked to Beck-Fahrner syndrome, BCL11A is associated with hereditary persistence of fetal hemoglobin and syndromic intellectual disability, SNAI2 is implicated in Waardenburg syndrome and neural crest differentiation, NEUROG2 is a proneural factor with roles in GABAergic interneuron fate, with PBX3, ATF7, and SKIL, regulating developmental patterning and transcriptional repression.

These results reveal convergent regulatory features across dopaminergic and GABAergic neurons. In both cell types, WT networks are characterized by transcription factors mainly associated with neuronal identity, developmental progression, and synaptic function, while in contrast, MECP2-networks are mainly characterized by chromatin remodelers, transcriptional repressors, and non-coding regulatory factors.

Despite cell-type–specific differences, both lineages converge on a shared set of master regulators. In the WT condition, MEIS2, NR2F1, NR2F2, and MECP2 were found in both dopaminergic and GABAergic systems. In the MECP2-condition, shared regulators included MEIS2, NR2F1, BCL11A and TCF4. Functional enrichment consistently linked these regulators to transcriptional control, chromatin remodeling, and neurodevelopmental disorders, including associations of TCF4 with Pitt-Hopkins syndrome.

Together, these findings indicate that MECP2 loss reorganizes transcriptional networks across neuronal subtypes, shifting them away from lineage-defining programs toward epigenetic regulators and repressors. The recurrent overlap of MEIS2, NR2F1, NR2F2, BCL11A and TCF4 highlights phenotype-relevant nodes that may underlie shared functional impairments in dopaminergic and GABAergic neuronal development.

### Results 4: Integration with known disease-association gene databases of Differentially expressed genes indicates key phenotypic agents

The Differentially expressed genes between WT and MECP2-were extracted as cell-type specific, afterwards genetic burden analysis was performed with ClinGen-SFARI and ClinGen-Genes4Epilepsy database (see Fig.4). For the ClinGen-SFARI comparisons, significant enrichment was observed for GABAergic (17/225 genes, P = 3.15 × 10−5, OR = 3.42), Maturing Neurons (40/714 genes, P = 6.03 × 10−7, OR = 2.55), and NPCs (43/1016 genes, P = 2.14 × 10−4, OR = 1.88). No significant enrichment was detected for Neuroblasts (1/33 genes, P = 0.55), Radial Glia (78/3537 genes, P = 0.80), or SPCs (20/575 genes, P = 0.06).

**Figure 4:**
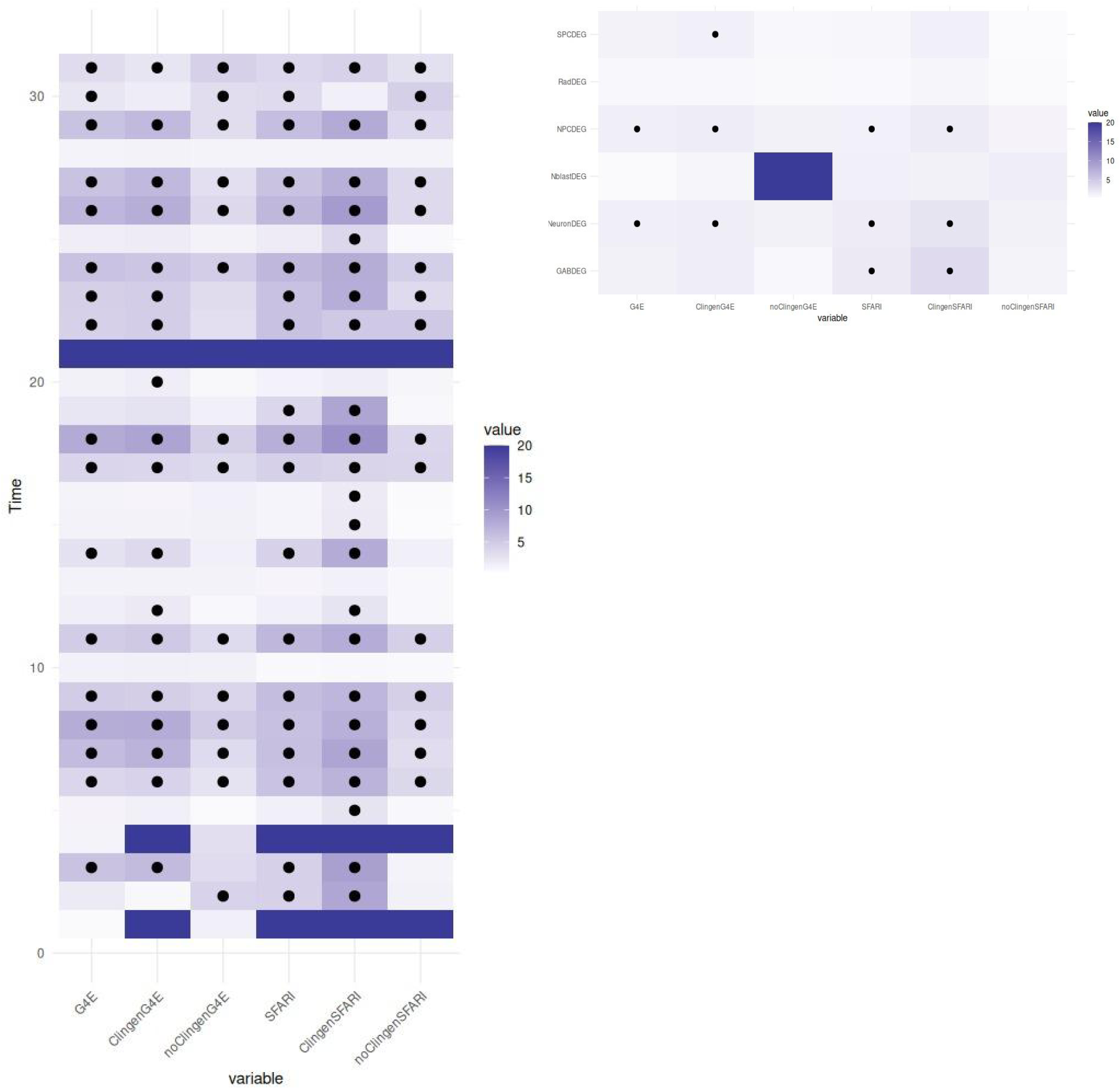
A) Heat map of enrichment in epilepsy-related genes (G4E), autism-related genes (SFARI), clinically validated autism and epilepsy-related genes (ClinGen), and not-ClinGen autism and epilepsy-related genes, divided by cell group Mark indicates significant enrichment, color indicates odd ratio with full band indicating NaN. B)Heat map of enrichment in G4E,SFARI, ClinGen, and not-ClinGen autism and epilepsy-related genes divided by Monocle Time.

For the ClinGen-Genes4Epilepsy comparisons, enrichment was significant for Maturing Neurons (38/714 genes, P = 3.37 × 10−3, OR = 1.66), NPCs (56/1016 genes, P = 1.70 × 10−4, OR = 1.75), and SPCs (27/575 genes, P = 4.86 × 10−2, OR = 1.44). No significant enrichment was detected for GABAergic (12/225 genes, P = 0.08), Neuroblasts (1/33 genes, P = 0.67), or Radial Glia (92/3537 genes, P = 0.99).

Notably, the strongest enrichments were consistently observed in Maturing Neurons and NPCs across both databases, while Radial Glia and Neuroblasts showed no enrichment which remains consistent with the boolean model performance, however while there is enrichment in validated genes related to disease along cell types, but not along detailed pseudotime groups.Comparison of the SFARI–G4Epi overlapping genes with the DopaMR and GabaMR marker lists provided revealed restricted but consistent intersections. In Maturing Neurons, four overlapping genes were also present in DopaMR (BCL11A, BCL11B, MEF2C, TCF4) and two in GabaMR (BCL11A, TCF4). In NPCs, three overlapping genes were also found in both DopaMR and GabaMR (BCL11A, NR2F1, TCF4). In SPCs, two overlaps were shared with DopaMR (MEF2C, NR2F1) and two with GabaMR (BPTF, NR2F1).

These results highlight a recurrent subset of transcriptional regulators (BCL11A, MEF2C, NR2F1, and TCF4) that not only drive the overlap between SFARI and G4Epi but are also represented in dopaminergic and GABAergic marker gene sets, underscoring their potential role as core mediators of neurodevelopmental risk across neuronal lineages.

### Results 5: Boolean model of proposed regulatory pathways

To explore the dynamic properties of the proposed master regulators and their progression during the development of the different cell types, we implemented dynamic modeling of the identified networks. This modeling utilized binarized gene expression data from developmental pathways of dopaminergic neurons, and gabaergic neurons with the combined MR sub-network translated into an influence graph utilizing the correlation between each pair of genes.

We started by modeling the system of master regulators in the Dopaminergic neurons (see Fig 5 A.), from this analysis we evaluated the attractor field of both conditions, calculating the amount of attractor states reachable from each time point, focusing on the groups that behaved differently between WT and MECP2-. Our first different group is Radial Glial cells in which we observed a significant loss of attractor states mapped (Fig 5 B.) with an extremely high convergence, when exploring the states and frequency between the WT condition (Fig 5 C.) and the Mecp2-condition(Fig 5 D.) we see no overlap on the states with higher convergence within the models with 4 states with higher convergence in the WT condition and 2 states with higher convergence in the MECP2-condition. Further exploring the behavior of the genes in the attractor states, both conditions stabilized NFIA and BCL11A, but MECP2⁻ introduced cyclic transitions involving ELK4 and RBPJ, absent from WT, which instead maintained broader static signatures with TCF4, SOX2, and VEZF1.

**Figure 5.**
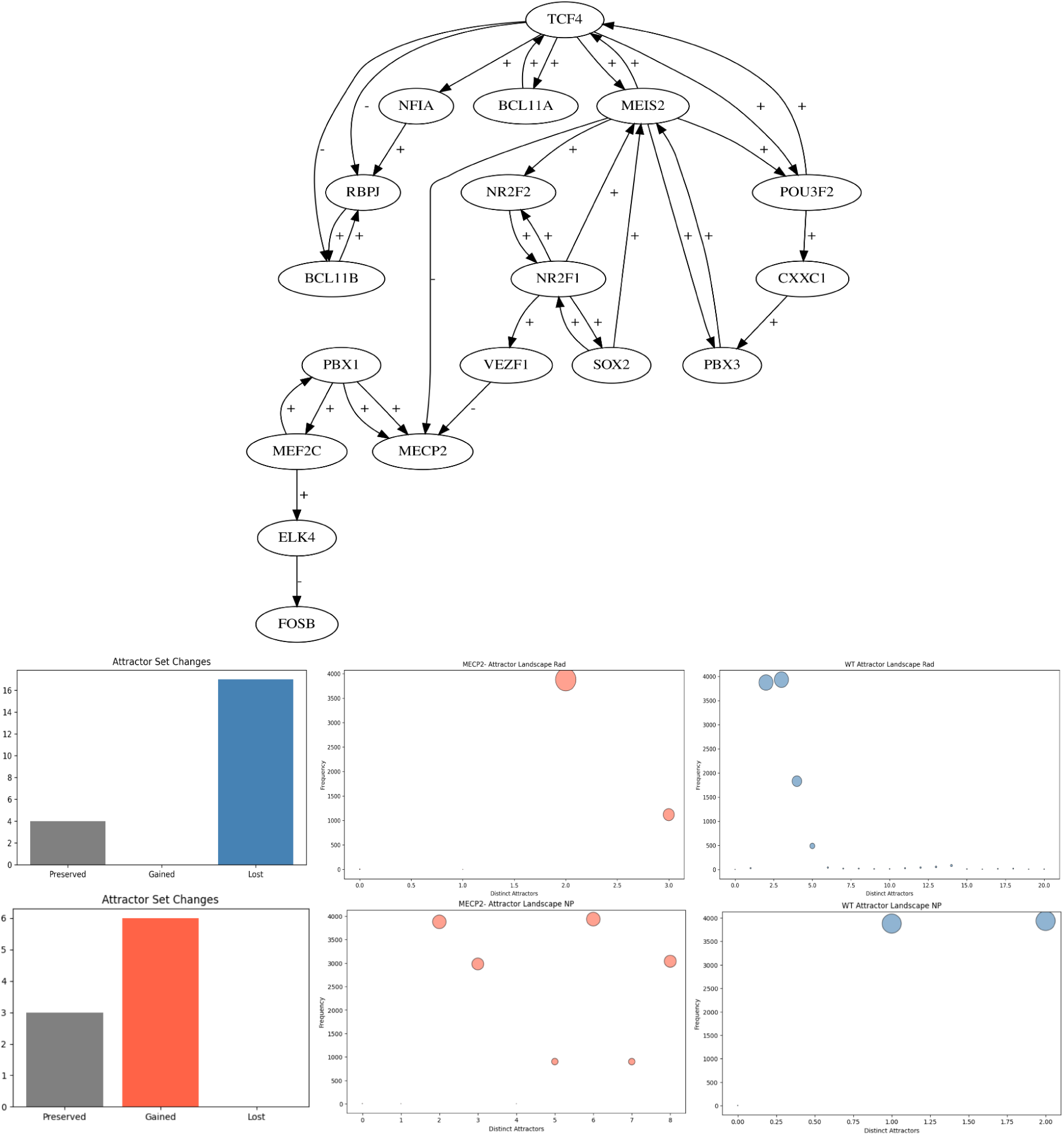
A) Influence Graph showcasing +/-relationship between Dopaminergic lineage Master Regulator network. B) Bargraph showcasing Attractor state changes in the MECP2-condition within Radial Glial Cells. C) Dotplot showcasing distincts attractors and their Frequency in the WT condition. D) Dotplot showcasing distincts attractors and their Frequency in the MECP2-condition. E) Bargraph showcasing Attractor state changes in the MECP2-condition within Neural Precursor Cells. F) Dotplot showcasing distincts attractors and their Frequency in the WT condition. G) Dotplot showcasing distincts attractors and their Frequency in the MECP2-condition.

In contrast Neural precursor cells we observed a significant gain of attractor states in the MECP2-condition mapped (Fig 5 E.) with little overlap between the modeled states, with 2 high convergence states in the Wt condition (Fig 5 F.) and 4 High convergence states in the MECP2-condition(Fig 5 G.). Exploring the behavior of the genes in the attractor states, Neural progenitors highlighted a major divergence between conditions with ELK4 - PBX3 cycles in MECP2-replacing the cyclic attractors with oscillations in RBPJ - BCL11B driven cycles of the WT Condition.

Other cell types did not showcase any variation in the attractor states frequency nor countwise. Further exploring the genes behavior within the model the attractor structures of WT and MECP2⁻ networks showed both conserved and divergent dynamics. Neuroblasts remained similar in their bifurcation between progenitor-like and neuron-primed states, though MECP2⁻ cycles emphasized CXXC1 and PBX3, whereas WT alternated between progenitor and differentiating attractors. Immature and mature neurons retained static expression of MEIS2, NR2F1/2, PBX3, and MEF2C across both conditions, but MECP2⁻ again favored ELK4- and SOX2-linked cycles, while WT cyclic activity prominently featured MECP2 and POU3F2. Finally, dopaminergic neurons stabilized into nearly identical static signatures with MEF2C, PBX1, and BCL11B, but WT additionally included NR2F2, while cyclic attractors in both conditions overlapped strongly around CXXC1, ELK4, and POU3F2.

Overall, static attractors defining progenitor and mature neuronal identities were conserved in MECP2⁻ cells, indicating that lineage specification remains largely intact. In contrast, cyclic attractor analysis revealed notable shifts: progenitors displayed increased attractor diversity, and oscillatory regulators shifted from RBPJ/MECP2 in WT to ELK4 in MECP2⁻. Terminal dopaminergic neurons showed incomplete activation of NR2F2 and continued cyclic transitions in CXXC1 and ELK4, highlighting altered regulatory dynamics despite preservation of core cell identities.

In the Gabaergic neurons model (see Fig 6 A.), from this analysis we evaluated the attractor field of both conditions, calculating the amount of attractor states reachable from each time point, focusing on the groups that behaved differently between WT and MECP2-this group showcased much more variety of states thus we focus in states in which the behavior outweighs the preserved behavior or in which the alteration occurs in state with high frequency. Our first different group is Neural progenitor gain of function in which we observed a significant gain of attractor states in the MECP2-condition mapped (Fig 6 B.) although there is significant overlap the differences occur within high frequency states between the modeled states, with 2 high convergence states in the Wt condition (Fig 5 F.) and 4 High convergence states in the MECP2-condition(Fig 5 G

**Figure 6).**
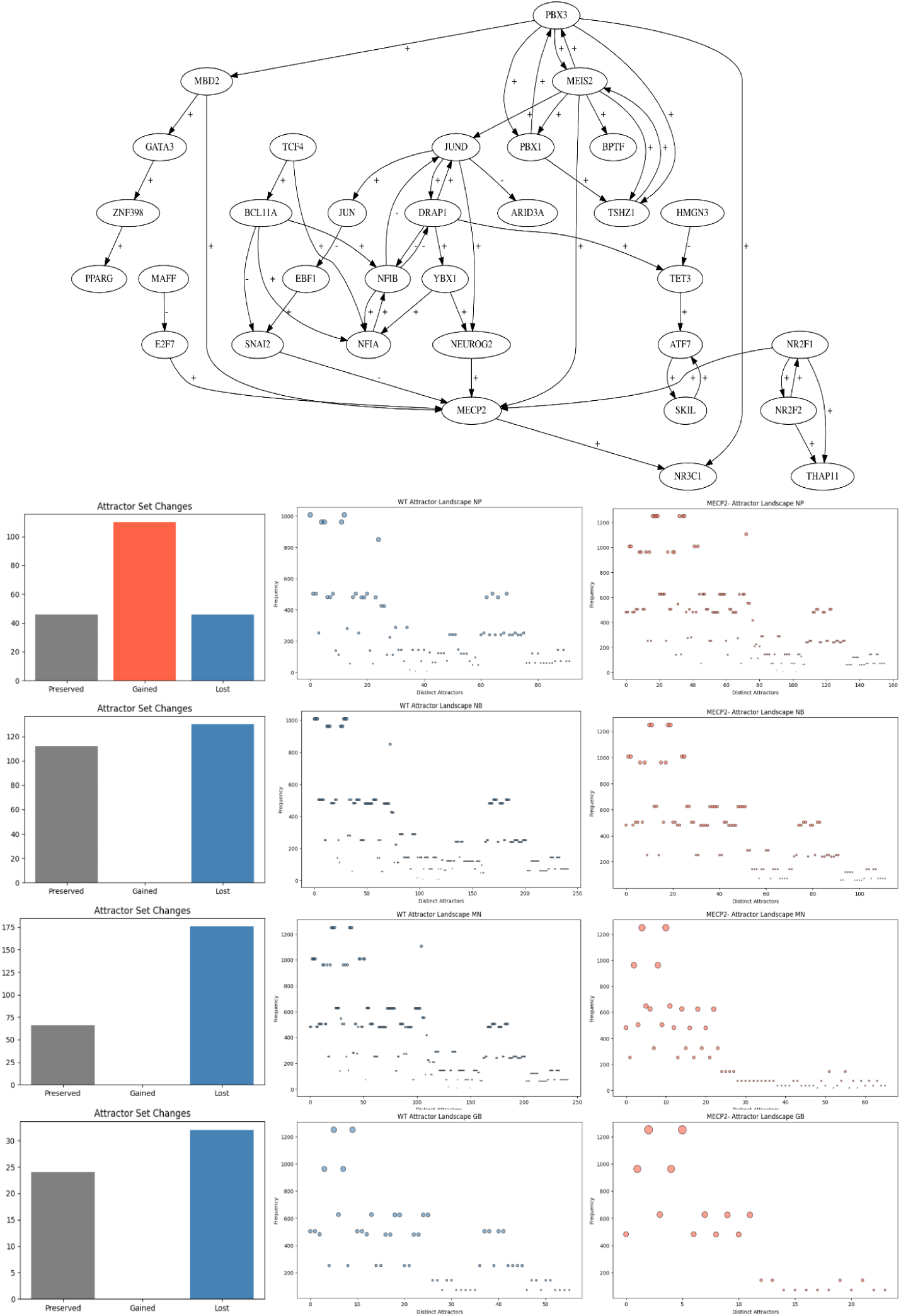
A) Influence Graph showcasing +/- relationship between Gabaergic lineage Master Regulator network. B) Bargraph showcasing Attractor state changes within Neural Precursors C-D) Dotplot showcasing distincts attractors and their Frequency in the WT condition and MECP2-condition. E) Bargraph showcasing Attractor state changes within Neuroblast F -G) Dotplot showcasing distincts attractors and their Frequency in the WT condition and the MECP2-condition.H) Bargraph showcasing Attractor state changes within Mature Neuron Cells I -J) Dotplot showcasing distincts attractors and their Frequency in the WT condition and the MECP2-condition. K) Bargraph showcasing Attractor state changes within Gabaergic neurons L -M) Dotplot showcasing distinct attractors and their Frequency in the WT condition and the MECP2-condition.

## Discussion

In this document we performed an in depth computational exploratory analysis of single cell organoid data in order to further unravel the complexities surrounding Rett syndrome phenotype and clinical implications this might have, previous literature describe the capability of cerebral organoids models to recapture developmental maturation of multiple neuron classes alongside their alteration by MeCP2 mutations[57,73], historically loss of MeCP2 function affects multiple neuronal developmental stages consistently between observation across models[74,75], with experimental evidence suggesting RTT to be a disorder of neuronal maturation and synapse formation, thus we looked to identify the key agents that could provide insight of this phenomena and their possible relationship to other conditions.

The key actors identified through our research landed on TCF4 and NF1R1 which consistently appeared across multiple network analyses. NF1R1 was identified in Module 3 of the WGCNA of Gabaergic neurons(figs.X), this also appeared as a WT condition master regulator in the Dopaminergic networks and a common master regulator between both condition in the Gabaergic networks (figs.X), in the clinically relevant differentially expressed genes, this gene is identified in the differentially expressed genes of Mature Neuron, Neural Precursor Cells, and Schwann cell precursors in the ClinGen-Genes4Epilepsy analysis, as well as in the differentially expressed genes of Gabaergic and Mature Neuron in the SFARI dataset(figs.X).

Regarding function NR2F1 is a transcription factor essential for multiple biological functions having been identified to work in regulating neural progenitor proliferation, neuronal migration, and cortical patterning [76]. It’s known for its dynamic expression, as the regulation of its multiple target genes gave a versatile yet contrasting role as previous research showed that during development it can both promote[76,77] or inhibit [76,78] proliferation, differentiation, and migration depending on developmental timing and cellular context in which it’s expressed. In neural progenitors, it regulates cell cycle dynamics to balance self-renewal and neurogenesis, while also influencing neuronal migration, axonal growth, and dendritic arborization[76,79]. Furthermore, NR2F1 controls progenitor identity, temporal competency, and contributes to establishing area-specific cortical identity[76,80]. Given the multifaceted nature of this gene its known that its dysfunction due to mutations cause Bosch–Boonstra–Schaaf optic atrophy syndrome (BBSOAS), as it results in problems on neuronal development and cortical organization, and their structural and functional abnormalities observed not only in this condition but with related neurodevelopmental disorders[76,81].

In the case TCF4, it was identified as a MECP2-condition master regulator in the Dopaminergic trajectory and a MECP2-condition master regulator in the Gabaergic trajectory (Fig.3), in the clinically relevant differentially expressed genes, this gene is identified in the differentially expressed genes of Mature Neuron and Neural Precursor Cells in the ClinGen-Genes4Epilepsy analysis, as well as in the differentially expressed genes of Gabaergic neurons, Mature Neuron and neural precursor in the ClinGen-SFARI dataset(Figs.4 A.).

Regarding function, the gene TCF4 is a transcription factor that plays a central role in nervous system development and function[82]. It is expressed during fetal brain development and persists throughout the adult forebrain and cerebellum, as well as in the oligodendrocytes of the spinal cord[82,83]. One of the main characteristics of TCF4 is its variety of isoforms as the functions of specific isoforms depend on which 5′-exon and internal exons that are included in the translated transcript, this also complicates the understanding of its molecular functions, as it’s through this mechanism that give it the ability to form heterodimers with other transcription factors,such as ATOH1, ASCL1, NEUROD1, and NEUROD2[84,85]. The more relevant functions for this case of study remain the differentiation of neuronal progenitor cells into neurons, maturation, neuronal migration and function, oligodendrocyte myelination and synaptic plasticity as we believe in its role in shaping both early neurodevelopmental processes and the maintenance of neuronal function throughout life[82,84,86].

Lineage-stage profiles serve as a proxy for regulatory activity in each lineage stage and enable an exploratory approach to conserved regulatory logic, functional convergence across models, thus, evaluating the behavior of the key actors detected across the models could elucidate highlight the behavior and changes that these genes have in the attractor sets with higher convergence defined by a significant basin size (>1000). This approach revealed that NF1R1 maintained a stable behavior as a static attractor in both the Dopaminergic neuron lineage and Gabaergic lineage for both WT and MECP2-specific models on the most stable attractors sets.Whereas TCF4 appeared to play a dynamic role in the WT models of both neural lineage models while in the MECP2-models it behaves as an static attractor in the most stable attractor sets, suggesting a that this behavior may be a compensatory role in the absence of MECP2.

Furthermore the lineage state models revealed an interesting pattern of behavior within the conditions, as regulatory dysregulation seems to be triggered at specific points of development to later converge in a standard behavior with these events occurring in key changes of differentiation potential within the neuron, this was more strongly seen in the radial glial cells and neural progenitors in the dopaminergic model and similarly in the neural progenitors,Neuroblast, Mature neurons, and Gabaergic neurons in the gabaergic model. We identified that the radial glial cell population corresponds to neural precursor glial cells, being within this cell type where we see the biggest loss of regulatory attractors between conditions, with this serving as a proxy of behavior and regulation, this strongly suggest that the dysregulation effects are first evidentiated in this stage as the radial glial cell transition to neural precursor, and furthermore down the line as we see Neuroblasts start the differentiation process we evidentiated a gain of attractor states, which would propose that some of the compensatory mechanisms within this regulatory pathway are activated to police this process.

In the context of RTT literary evidence demonstrates that the phenotype alterations of the syndrome are present at the earliest stages of brain development due to the pleiotropic effects of MeCP2[87–89], the impact of this are marked in developmental stages that define neurogenesis, migration, and patterning[23,87,89], which correlates with the master regulator behavior observed in the dynamic models.

Particularly notable is the rhythm of TCF4 in the MECP2-models, as previous literature with substantial experimental data has demonstrated the important role of the TCF4 transcription factor in the development of the nervous system with functional anomalies being known to drive the development of Pitt–Hopkins syndrome (PTHS)[82,84]. However, is in this context that an interesting paradigm arrives which is that in PTHS condition MECP2 results in behavioral phenotypic rescue in *Tcf4^+/-^* mice, with the experimental data indicating that this is results from differential glial cell development mediated by MeCP2 overexpression[88]. Correlating back to the behavior proposed by our results, with the pattern evidentiated within them, we suggest that this phenotypic rescue is compensatory in nature for both MECP2 and TCF4, furthermore we propose that the key to this compensatory response could be located in this changes of potentiality between the precursors cells, and could be the key to understand not only RTT but also PTHS.

Despite this however we remain at lost of the nature of NRF1 within the system, within the dopaminergic models NR2F1 seems to play an excitatory role, as its targets identified showcase a positive correlation in both WT and MECP2-in both lineage models, in further insight Enrichment of the common differentially expressed genes within the relevant cell types suggest that this system might be modeled by the Mark-Erk pathway[90]. Analysis of literature regarding this relationship showcases that there is experimental precedence of the immune model of neural development to impact RTT[23,74,90,91] with recent publications showcasing that rescue of the microglial proinflammatory cytokine production a*fter symptom onset* showing promise in improving neurobehavioral impairments along with sleep pattern and epileptiform activity[92] burden with precedence of the immune model also having a critical impact in several broader neurodevelopmental disorders further expanding on the impact of this results[93,94].

However further experimental research of the points presented by this analysis would be required to cement the finding in this study, but we remained satisfied as we propose a novel angle for Rett syndrome research that could open new approaches for therapeutic intervention and further our understanding of this condition.

### Conclusions

This exploratory analysis revealed cellular subtypes within the organoids, showcasing differences in the distribution of neurons, pseudo time calculations enabling the reconstruction of developmental sequences, providing a comprehensive understanding of the disrupted developmental pathways of Rett syndrome.

Network analysis identified master regulators responsible for modulating expression in GABAergic neurons lineage as well as Dopaminergic neurons lineage. Identifying a system with MR exclusive to MECP2+ neurons and MR exclusive to MECP2-neurons, co-regulating each other providing insight within Rett syndrome’s compensatory mechanisms with dynamic

Our study provides an insight that pinpoints the cellular stages on which the regulation and compensatory mechanism activate and regulate Rett syndrome. We identified 19 Master regulators for the Dopaminergic developmental trajectory, as well as 34 Master regulator genes for the gabaergic developmental trajectory. Dynamic Boolean modeling of this systems showcased both attractor landscape as a proxy for regulatory activity in each lineage stage providing a comprehensive understanding of the disrupted developmental pathways of Rett syndrome, highlighting the transitional states of potential within maturation trajectories as the key point of divergence in regulation for Rett syndrome.

After complementing with enrichment and clinical relevant variant analysis, we identify the key actors in this system as NR2F1 and TCF4, with TCF4 indicating a symmetrical compensatory relationship with MeCP2, and NR2F1 a possible link with wider developmental conditions, this concluded with highlighting the possibility of regulation in this condition being affected by the MAPK-ERK pathway of transcriptional regulation, corroborated by historical Rett syndrome reported experimental research.

This provides novel paradigms to both possible new therapeutic avenues as well as key agents for targeted experimental research that can not only provide a better understanding of Rett syndrome but of neurodevelopmental conditions as a whole, highlighting the role of the Immune modulation of neural regulation during development.

## Acknowledgement

Powered@NLHPC supercomputing infrastructure of the NLHPC (ECM-02)

